# Knowledge is power: Contingency instruction promotes threat extinction in high intolerance of uncertainty individuals

**DOI:** 10.1101/470724

**Authors:** Jayne Morriss, Carien M. van Reekum

## Abstract

Extinction-resistant threat is considered to be a central feature of pathological anxiety. Reduced threat extinction is observed in individuals with high intolerance of uncertainty (IU). Here we sought to determine whether contingency instructions could alter the course threat extinction for individuals high in IU. We tested this hypothesis in two identical experiments (Exp 1 *n* = 60, Exp 2 *n* = 82) where we recorded electrodermal activity during threat acquisition with partial reinforcement, and extinction. Participants were split into groups based on extinction instructions (instructed, uninstructed) and IU score (low, high). All groups displayed larger skin conductance responses to learned threat versus safety cues during threat acquisition, indicative of threat conditioning. In both experiments, only the uninstructed high IU groups displayed larger skin conductance responses to the learned threat versus safety cue during threat extinction. These findings suggest that uncertain threat during extinction maintains conditioned responding in individuals high in IU.

## Introduction

The ability to discriminate between threat and safety is crucial for maintaining health and wellbeing. Through threat conditioning, an organism can associate neutral cues (conditioned stimulus, e.g. a visual stimulus such as a shape) with aversive outcomes (unconditioned stimulus, e.g. shock, loud tone). Repeated presentations of a neutral cue with an aversive outcome can result in threat responding to the conditioned cue (conditioned response). This learned association can also be extinguished by repeatedly presenting the conditioned cue without the aversive outcome (LeDoux, 1998; Myers & Davis, 2007). The reduction in reactivity observed to the conditioned cue over time is thought to reflect changes in contingency beliefs e.g. the threat becomes safe (Hofmann, 2008).

Notably, in anxiety and stress disorders, physiological responses are exaggerated and sustained to cues that no longer signal threat, suggesting impaired threat extinction (Blechert, Michael, Vriends, Margraf, & Wilhelm, 2007; Michael, Blechert, Vriends, Margraf, & Wilhelm, 2007; Milad et al., 2008; Milad et al., 2009). Disrupted threat extinction in anxious individuals is likely maintained through greater expectations of threat, also known as threat expectancy biases (Craske, Treanor, Conway, Zbozinek, & Vervliet, 2014; Hofmann, 2008). One potential factor that may prevent or prolong threat extinction is uncertainty surrounding the contingency change due to the omission of the US. Uncertainty has been identified as an important facet of anxiety and stress disorders (Carleton, 2016a, 2016b; Dugas, Buhr, & Ladouceur, 2004; Grupe & Nitschke, 2013). Despite this, only recently has the role of individual differences in Intolerance of Uncertainty (IU), a tendency to find uncertainty anxiety provoking, been examined in relation to threat extinction (Dunsmoor, Campese, Ceceli, LeDoux, & Phelps, 2015; Lucas, Luck, & Lipp, 2018; Morriss, Christakou, & van Reekum, 2015, 2016; Morriss, Macdonald, & van Reekum, 2016). Previous work has shown that high IU is associated with greater skin conductance responding to learned threat versus safety cues during extinction (Morriss, Christakou, & van Reekum, 2015, 2016; Morriss, Macdonald, & van Reekum, 2016). Furthermore, individuals high in IU are more prone to spontaneous recovery of learned threat during next day extinction (Dunsmoor, Campese, Ceceli, LeDoux, & Phelps, 2015; Lucas, Luck, & Lipp, 2018). Overall, these results suggest that individual differences in IU modulate threat expectancy biases during threat extinction.

Questions remain on how IU modulates threat expectancy biases during threat extinction. For example, is it the uncertainty surrounding the omission of the US that prolongs threat extinction learning? A way to address this question is to give individuals high in IU more information and hence reduce uncertainty about the US omission to observe whether this promotes threat extinction. Previous research has demonstrated that providing instructions about threat and safety contingencies speeds up the course of threat extinction (Javanbakht et al., 2017; Koenig & Henriksen, 2005; Sevenster, Beckers, & Kindt, 2012). The effect of instruction on threat extinction is robust and has been found using a variety of conditioning designs with different psychophysiological measures (but not fear relevant conditioned stimuli such as snakes) (Luck & Lipp, 2016). However, to date there is a dearth of research on the effect of instructed threat extinction in subclinical and clinical anxiety, or in individuals high in IU. Given the significant role of uncertainty in anxiety (Carleton, 2016a, 2016b; Grupe & Nitschke, 2013) and that current exposure therapies are based on associative learning principles (Craske, Treanor, Conway, Zbozinek, & Vervliet, 2014), examining the effect of instructed threat extinction on individuals high in IU may reveal vital information relevant to IU-related threat expectancy biases. Furthermore, such examinations may open avenues for future threat extinction research and exposure-based treatments for anxiety and stress disorders.

In two identical experiments we used an instructed threat extinction paradigm, in order to assess the relationship between individual differences in self-reported IU and threat expectancy biases. We measured skin conductance responses (SCR) and expectancy ratings whilst participants underwent threat acquisition and extinction phases. We used an aversive sound as an unconditioned stimulus and visual shape stimuli as conditioned stimuli, similar to previous conditioning research including our own (Morriss, Christakou, & van Reekum, 2015, 2016; Morriss, Macdonald, & van Reekum, 2016; Neumann, Waters, & Westbury, 2008). We used a 50% reinforcement rate during acquisition to sustain conditioning during extinction (Leonard, 1975; Livneh & Paz, 2012). We had four groups of participants: low IU uninstructed; low IU instructed; high IU uninstructed; high IU instructed. Prior to threat extinction, participants in the instruction groups were presented with the threat and safety contingencies, whilst the uninstructed groups received no information about the change in contingencies.

We hypothesised that during threat acquisition, skin conductance responding and expectancy ratings would be higher to the learned threat versus safety cues. Based on previous research, we predicted that only the high IU uninstructed group would exhibit greater skin conductance and expectancy ratings to the learned threat versus safety cues during extinction (Morriss, Christakou, & van Reekum, 2015, 2016; Morriss, Macdonald, & van Reekum, 2016). Furthermore, we predicted that the other three groups would be capable of threat extinction, albeit for different reasons. We predicted that low IU individuals would extinguish regardless of instruction, as they don’t find uncertainty aversive. In addition, we predicted the high IU instructed group to extinguish, as the instructions would reduce uncertainty about the US omission. In line with our previous work (for discussion see Morriss, Christakou & van Reekum, 2016) we tested the specificity of IU effects by controlling for trait anxiety, assessed by the commonly used Spielberger State-Trait Anxiety Inventory, Trait Version (STAI) (Spielberger, Gorsuch, Lushene, Vagg, & Jacobs, 1983).

## Experiment 1: Method

### Participants

Sixty volunteers (*M* age = 23.56, *SD* age = 4.58; 33 females and 27 males) took part in the study. All participants had normal or corrected to normal vision. Participants provided written informed consent and received £5 for their participation. Advertisements and word of mouth were used to recruit participants from the University of Reading and local area. Participants were recruited if they were between 18-40 years of age. No other exclusion criteria were used. One participant withdrew from the experiment and one participant had incomplete questionnaire data, leaving fifty-eight participants with usable data. The procedure was approved by the University of Reading Research Ethics Committee.

### Procedure

Prior to arrival at the laboratory, participants were emailed two questionnaires to assess their anxious disposition. Group allocation was based on a median split. Depending upon whether participants scored high (above average, <65) or low (below average, >65) on the IU questionnaire (Freeston, Rhéaume, Letarte, Dugas, & Ladouceur, 1994) participants were allocated to an instructed or uninstructed condition, thus creating four groups: low IU instructed (n = 14); low IU uninstructed (n = 15); high IU uninstructed (n =13); high IU instructed (n = 16). Different researchers were responsible for participant grouping and data collection to allow the interacting researcher to remain blind to participants’ IU score.

On the day of the experiment participants arrived at the laboratory and were informed on the experimental procedures. Firstly, participants were seated in the testing booth and asked to complete and sign a consent form as an agreement to take part in the study. Secondly, physiological sensors were attached to the participants’ non-dominant hand. The conditioning task (see “Conditioning task” below for details) was presented on a computer, whilst skin conductance, interbeat interval and behavioural ratings were recorded. Participants were instructed to: (1) maintain attention to the task by looking at the coloured squares and listening to the sounds, which may be unpleasant, (2) respond to the expectancy rating scales that followed each block of trials, using number keys on the keyboard with their dominant hand and (3) to stay as still as possible. The experiment took approximately 30 minutes in total.

### Conditioning task

The conditioning task was designed using E-Prime 2.0 software (Psychology Software Tools Ltd, Pittsburgh, PA). Visual stimuli were presented at a 60 Hz refresh rate on an 800 x 600 pixel computer screen. Participants sat approximately 60 cm from the screen. Visual stimuli were blue and yellow squares with 183 × 183 pixel dimensions that resulted in a visual angle of 5.78° × 9.73°. The aversive sound stimulus was presented through headphones. The sound consisted of a fear inducing female scream used in our previous experiments (Morriss et al., 2015; Morriss et al., 2016). The volume of the sound was standardized across participants by using fixed volume settings on the presentation computer and was verified by an audiometer prior to each session (90 dB).

The task comprised of two learning phases: acquisition and extinction. Both acquisition and extinction consisted of two blocks each. In acquisition, one of the coloured squares (blue or yellow) was paired with the aversive 90 dB sound 50% of the time (CS+), whilst the other square (yellow or blue) was presented alone (CS-). The 50% pairing rate was designed to maximize the unpredictability of the CS+ / US contingency. Prior to extinction, participants in the instruction condition were presented the following statement: “From now on the blue/yellow square (i.e. CS+) will no longer be paired with an aversive sound. The yellow/blue square (i.e. CS-) will continue to be presented alone without any sound”. Participants were asked to confirm they understood this statement through intercom before the extinction phase began. Participants in the uninstructed condition were not presented with the statement. During extinction, both the blue and yellow squares were presented in the absence of the US; this was true for both instructed and uninstructed conditions.

The acquisition phase consisted of 24 trials (6 CS+ paired, 6 CS+ unpaired, 12 CS-) and the extinction phase 32 trials (16 CS+ unpaired, 16 CS-). Experimental trials were pseudo-randomised such that the first trial of acquisition was always paired and then after all trial types were randomly presented. Conditioning contingencies were counterbalanced, with half of participants receiving the blue square paired with the US and the other half of participants receiving the yellow square paired with the US. The coloured squares were presented for a total of 4000 ms. The aversive sound lasted for 1000 ms, which coterminated with the reinforced CS+’s. Subsequently, a blank screen was presented for 6000 – 8800 ms.

At the end of each block, participants were asked to rate how much they expected the blue square and yellow square to be followed by the sound stimulus, where the scale ranged from 1 (“Don’t Expect”) to 9 (“Do Expect”). Four other 9-point Likert scales were presented at the end of the experiment. Participants were asked to rate: (1) the valence and (2) arousal of the sound stimulus, as well as (3) the valence and (4) arousal of the unpredictability of the sound. The scales ranged from 1 (Valence: very negative; Arousal: calm) to 9 (Valence: very positive; Arousal: excited).

### Questionnaires

To assess anxious disposition, we administered the STAI (Spielberger, Gorsuch, Lushene, Vagg, & Jacobs, 1983) and IU questionnaires (Freeston, Rhéaume, Letarte, Dugas, & Ladouceur, 1994). The IU measure consists of 27 items. Items include “Uncertainty makes me uneasy, anxious, or stressed” and “I must get away from all uncertain situations”. Similar distributions and internal reliability of scores were found for the anxiety measures, STAI (*M* = 41.81; *SD* = 10.64; range = 24-60; *α* = .92), IU (*M* = 67.53; *SD* = 17.41; range = 29-100; *α* = .92). The instructed and uninstructed groups were matched on IU: low IU uninstructed (*M* = 52.14; *SD* = 7.32); low IU instructed (*M* = 53.8; *SD* = 10.13); high IU uninstructed (*M* = 80.94; *SD* = 10.54); high IU instructed (*M* = 83.46; *SD* = 9.79).

### Behavioural data scoring

Rating data were reduced for each participant by calculating their average responses for each experimental condition using the E-Data Aid tool in E-Prime (Psychology Software Tools Ltd, Pittsburgh, PA).

### Physiological acquisition and scoring

Physiological recordings were obtained using AD Instruments (AD Instruments Ltd, Chalgrove, Oxfordshire) hardware and software. Electrodermal activity was measured with dry MLT116F silver/silver chloride bipolar finger electrodes that were attached to the distal phalanges of the index and middle fingers of the non-dominant hand. A low constant-voltage AC excitation of 22 mV_rms_ at 75 Hz was passed through the electrodes, which were connected to a ML116 GSR Amp, and converted to DC before being digitized and stored. Interbeat Interval (IBI) was measured using a MLT1010 Electric Pulse Transducer, which was connected to the participant’s distal phalange of the ring finger. An ML138 Bio Amp connected to an ML870 PowerLab Unit Model 8/30 amplified the skin conductance and IBI signals, which were digitized through a 16-bit A/D converter at 1000 Hz. IBI signal was only used to identify movement artefacts and was not analysed. The electrodermal signal was converted from volts to microSiemens using AD Instruments software (AD Instruments Ltd, Chalgrove, Oxfordshire).

CS+ unpaired and CS− trials were included in the analysis, but CS+ paired trials were discarded to avoid sound confounds. Skin conductance responses (SCR) were scored when there was an increase of skin conductance level exceeding 0.03 microSiemens (Dawson, Schell, & Filion, 2000). The amplitude of each response was scored as the difference between the onset and the maximum deflection prior to the signal flattening out or decreasing. SCR onsets and respective peaks were counted if the SCR onset was within 0.5-3.5 seconds (CS response) following CS onset (Morriss, Chapman, Tomlinson, & van Reekum, 2018). Trials with no discernible SCRs were scored as zero (percentage of CS+ unpaired and CS− trials scored as zero during: Acquisition, 46%; Extinction, 56.0%). SCR magnitudes were square root transformed to reduce skew and were z-scored to control for interindividual differences in skin conductance responsiveness (Ben-Shakhar, 1985). SCR magnitudes were calculated from remaining trials by averaging SCR square-root-transformed values and zeros for each condition. We defined non-responders as those who responded to 10% or less of the CS+ unpaired and CS− trials. From this we identified 1 non-responder from the high IU uninstructed group, who we removed from the subsequent analyses, leaving fifty-seven participants with usable SCR data.

### Ratings and SCR magnitude analysis

We conducted separate within-between repeated measures ANCOVA’s on ratings and SCR during threat acquisition and extinction. For Acquisition, we conducted a 2 Condition (CS+, CS−) x 4 Group (high IU instructed, high IU uninstructed, low IU instructed and low IU uninstructed) x STAI. For extinction, we conducted a 2 Condition (CS+, CS−) x Time (Early, Late) x 4 Group (high IU instructed, high IU uninstructed, low IU instructed and low IU uninstructed) x STAI. We included STAI as a covariate to assess the specificity of IU.

## Experiment 1: Results

### Ratings

Participants rated the sound stimulus as aversive (*M* = 2.34, *SD* = 1.2, range 1-7, where 1 = very negative and 9 = very positive) and arousing (*M* = 6.80, *SD* = 1.6, range 2-9 where 1 = calm and 9 = excited).

For the expectancy ratings, during acquisition participants reported greater expectancy of the sound with the CS+, compared to CS− [Stimulus: *F*(1, 53) = 42.202, *p* < .001, □^2^ =.44] (for descriptive statistics see Table 1). No other significant interactions with IU group or STAI were found for the ratings during acquisition, max *F* =1.406.

**Table 1.** Experiment 1 summary of means (SD) for each dependent measure as a function of stimulus (CS+ and CS−), separately for acquisition, early extinction and late extinction.

| Measure | Acquisition |  | Early Extinction |  | Late Extinction |  |
| --- | --- | --- | --- | --- | --- | --- |
|  | CS+ | CS- | CS+ | CS- | CS+ | CS- |
| Square root transformed and z-scored SCR magnitude ($\sqrt{\mu\text{S}}$) | 0.28<br>(0.43) | 0.01<br>(0.28) | 0.00<br>(0.38) | -0.09<br>(0.38) | -0.04<br>(0.36) | -0.09<br>(0.39) |
| Expectancy rating (1-9) | 4.30<br>(1.14) | 2.87<br>(1.70) | 7.02<br>(1.20) | 1.81<br>(1.48) | 2.64<br>(2.00) | 1.82<br>(1.87) |

Note: SCR magnitude ( $\sqrt{\mu\text{S}}$ ), square root transformed and z-scored skin conductance magnitude measured in microSiemens.

During extinction, participants reported greater expectancy of the sound with the CS+, compared to CS− [Stimulus: *F*(1, 53) = 104.445, *p* < .001, □^2^ =.66]. The expectancy ratings dropped over time [Time: *F*(1, 53) = 104.445, *p* < .001, □^2^ =.66; Stimulus x Time: *F*(1, 53) = 206.779, *p* < .001, □^2^ =.79]. Follow-up pairwise comparisons revealed that the expectancy rating of the sound with the CS+ dropped significantly from early to late extinction, p < .001. However, the expectancy rating of the CS− with the sound remained low and did not change with time, p = .906. Unexpectedly there was an interaction with STAI [Stimulus x Time x STAI: *F*(1, 53) = 4.234, *p* = .045, □^2^ =.07], carried by individuals high in trait anxiety who showed a reduction in expectancy of the sound with the CS− from early (*M* = 3.10, *SE* = .396) to late (*M* = 1.66, *SE* = .391) extinction, *p* < .001, whereas individuals low in trait anxiety showed similar ratings of expectancy to the sound with the CS− across early (*M* = 2.19, *SE* = .398) to late (*M* = 2.03, *SE* = .393) extinction, *p* = .273. No other significant interactions with IU group or STAI were found for the ratings during extinction, max *F* = 1.502.

### SCR magnitude

During acquisition participants displayed greater SCR magnitude to the CS+, compared to CS− [Stimulus: *F*(1, 52) = 18.626, *p* < .001, □^2^ =.26] (for descriptive statistics see Table 1). No significant interactions with IU group (or STAI) were found for SCR magnitude during acquisition, max *F* =1.083.

During extinction, only the uninstructed high IU group displayed larger SCR magnitude to the CS+ vs. CS, *p* = .002 [Stimulus x IU group: *F*(1, 52) = 3.047, *p* = .037, □^2^ =.15] (see Figure 1). The other 3 remaining groups displayed no significant differences between CS+ vs. CS, *p*’s > .5^1^. The SCR magnitude for the CS+ was significantly larger for the uninstructed high IU group, vs. the uninstructed low IU group, *p* = .030 and the instructed high IU group, *p* =.038. In addition, the SCR magnitude for the CS− was significantly reduced for the uninstructed high IU group, vs. the uninstructed low IU group, *p* = .047 and the instructed low IU group, *p* =.030. All other multiple comparisons from this interaction were above *p* >.05. No other significant interactions with Time, IU group or STAI were found for SCR magnitude during extinction, max *F* =1.458.

**Fig 1.**
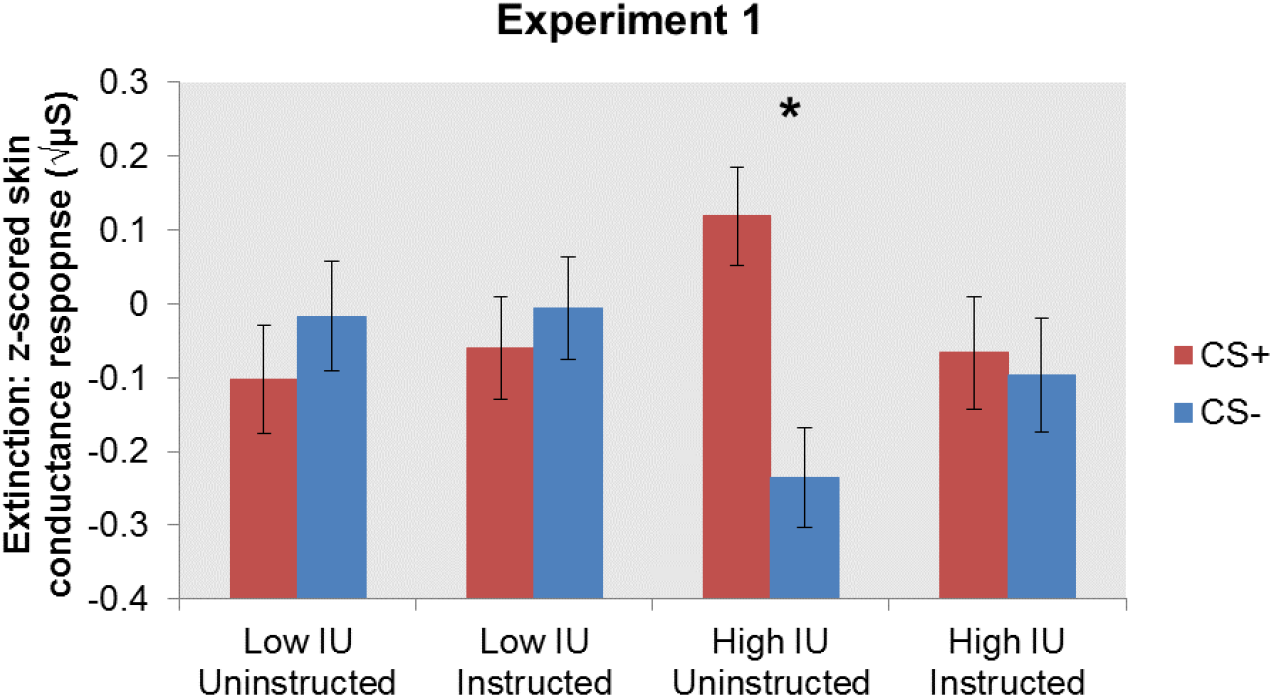
Experiment 1 SCR magnitude results for IU group (controlling for STAI) during threat extinction. Only the high IU uninstructed group were found to show differential skin conductance responding to the CS+ versus CS− cue during threat extinction. Bars represent standard error. Square root transformed and z-scored SCR magnitude (μS), skin conductance magnitude measured in microSiemens. Note that the z-scoring was performed within-subjects, across both phases, thus explaining the negative values for most conditions.

## Experiment 1: Conclusion

For experiment 1 we observed typical profiles of acquisition, where larger SCR magnitudes and expectancy ratings were found for the CS+ vs. CS−, across all groups. In addition, for extinction we observed a reduction in expectancy ratings of the sound for the CS+ vs. CS−. During extinction, only the uninstructed high IU group displayed larger SCR magnitudes to the CS+ vs. CS. The other three groups showed no differential SCR magnitudes between the CS+ vs. CS, indicative of extinction.

The lack of extinction in the uninstructed high IU group partially replicated our previous IU and uninstructed extinction research (Morriss, Christakou, & Van Reekum, 2015, 2016). We observed no IU differences on the ratings. However, we did observe an effect of STAI on the ratings during extinction.

## Experiment 2: Method

The method was identical to experiment 1, except for details provided below.

### Participants

Eighty-two volunteers (*M* age = 24.65, *SD* age = 4.30; 57 females, 24 males, 1 missing information for sex) took part in the study. We based our sample size on a power analysis using the effect size (.15) from the Stimulus x IU group interaction for SCR magnitude in experiment 1. The following parameters were used for a repeated measures within-between interaction design: effect size *f* = 0.15, α error probability = 0.05, Power (1-β error probability) = 0.7, number of groups = 4 (low IU uninstructed, low IU instructed, high IU uninstructed, high IU instructed). The total sample size suggested was 76 (19 per group). We oversampled due to expected participant attrition from non-responding in SCR magnitude. One participant withdrew from the experiment, leaving eighty-one participants with usable data.

### Procedure

The procedure was identical to experiment 1, except that the questionnaires were completed on a computer on the day of testing. Participants were allocated to one of four groups based on their IU score (the cut-off was identical to Experiment 1): low IU uninstructed (n = 21); low IU instructed (n = 22); high IU uninstructed (n =19); high IU instructed (n = 19). As in Experiment 1, different researchers were responsible for participant grouping and data collection to allow the interacting researcher to remain blind to participants’ IU score.

### Questionnaires

Distributions and internal reliability of scores were similar to those found in Experiment 1 for the anxiety measures, STAI (*M* = 43.80; *SD* = 9.31; range = 26-68; *α* = .89), IU (*M* = 65.96; *SD* = 18.07; range = 33-100 *α* = .92). The instructed and uninstructed groups were matched on IU: low IU uninstructed (*M* = 50.66; *SD* = 8.69); low IU instructed (*M* = 52.63; *SD* = 9.51); high IU uninstructed (*M* = 82.00; *SD* = 10.60); high IU instructed (*M* = 82.26; *SD* = 10.42).

### Physiological acquisition and scoring

Percentage of CS+ unpaired and CS− trials scored as zero during: Acquisition, 45%; Extinction, 51%. We identified 2 non-responders, one from the uninstructed low group and one from the uninstructed high group, who we removed from the subsequent analyses, leaving seventy-nine participants with usable SCR data.

## Experiment 2: Results

### Ratings

In general, participants rated the sound stimulus as aversive (*M* = 2.22, *SD* = 1.43, range 1-7, where 1 = very negative and 9 = very positive) and arousing (*M* = 6.93, *SD* = 1.73, range 2-9 where 1 = calm and 9 = excited).

For the expectancy ratings, during acquisition participants reported greater expectancy of the sound with the CS+, compared to CS− [Stimulus: *F*(1, 76) = 94.734, *p* < .001, □^2^ =.55] (for descriptive statistics see Table 2). No significant interactions with IU group or STAI were found for the ratings during acquisition, max *F* =1.040.

**Table 2.** Experiment 2 summary of means (SD) for each dependent measure as a function of stimulus (CS+ and CS−), separately for acquisition, early extinction and late extinction.

| Measure | Acquisition |  | Early Extinction |  | Late Extinction |  |
| --- | --- | --- | --- | --- | --- | --- |
|  | CS+ | CS- | CS+ | CS- | CS+ | CS- |
| Square root transformed and z-scored SCR magnitude ($\sqrt{\mu\text{S}}$) | 0.12<br>(0.48) | -0.01<br>(0.29) | 0.04<br>(0.38) | -0.08<br>(0.33) | 0.02<br>(0.35) | -0.06<br>(0.44) |
| Expectancy rating (1-9) | 6.49<br>(1.98) | 2.72<br>(2.12) | 4.41<br>(2.63) | 2.83<br>(2.68) | 3.30<br>(2.40) | 2.53<br>(2.46) |

Note: SCR magnitude ( $\sqrt{\mu\text{S}}$ ), square root transformed and z-scored skin conductance magnitude measured in microSiemens.

During extinction, participants reported greater expectancy of the sound with the CS+, compared to CS− [Stimulus: *F*(1, 76) = 23.683, *p* < .001, □^2^ =.23]. Participants expectancy ratings dropped over time [Time: *F*(1, 76) = 19.743, *p* < .001, □^2^ =.20; Stimulus x Time: *F*(1, 76) = 11.350, *p* < .001, □^2^ =.13]. Follow-up pairwise comparisons showed that the expectancy rating of the sound with the CS+ significantly reduced across early to late extinction, *p* < .001. In addition, there was a trend for the expectancy rating of the CS− with the sound to drop across early to late extinction time, *p* = .052. No other significant interactions with IU group or STAI were found for the ratings during extinction, max *F* = 1.996.

### SCR magnitude

During acquisition participants displayed greater SCR magnitude to the CS+, compared to CS− at trend [Stimulus: *F*(1, 74) = 3.250, *p* = .076, □^2^ =.04] (for descriptive statistics see Table 2). No significant interactions with IU group or STAI were found for SCR magnitude during acquisition, max *F* = .801.

During extinction, participants displayed greater SCR magnitude to the CS+ vs. CS− [Stimulus: *F*(1, 74) = 5.655, *p* = .020, □^2^ =.07]. This main effect was likely driven by the uninstructed high IU group, as this was the only group to display larger SCR magnitude to the CS+ vs. CS, *p* = .005 [Stimulus x IU group: *F*(1, 74) = 2.948, *p* = .038, □^2^ =.10] (see Figure 2). The other 3 remaining groups displayed no significant differences for SCR magnitude between the CS+ and CS, *p*’s > .19^2^. The SCR magnitude for the CS+ was significantly larger for the uninstructed high IU group, vs. the instructed high IU group, *p* =.003. In addition, the magnitude of the response to the CS+ was significantly larger for both low IU groups, compared to the high IU instructed group, p’s < .036. Furthermore, the SCR magnitude for the CS− was significantly reduced for the uninstructed high IU group, vs. the instructed high IU group, *p* =.040. No other significant interactions with Time, IU group or STAI were found for SCR magnitude during extinction, max *F* =.711.

**Fig 2.**
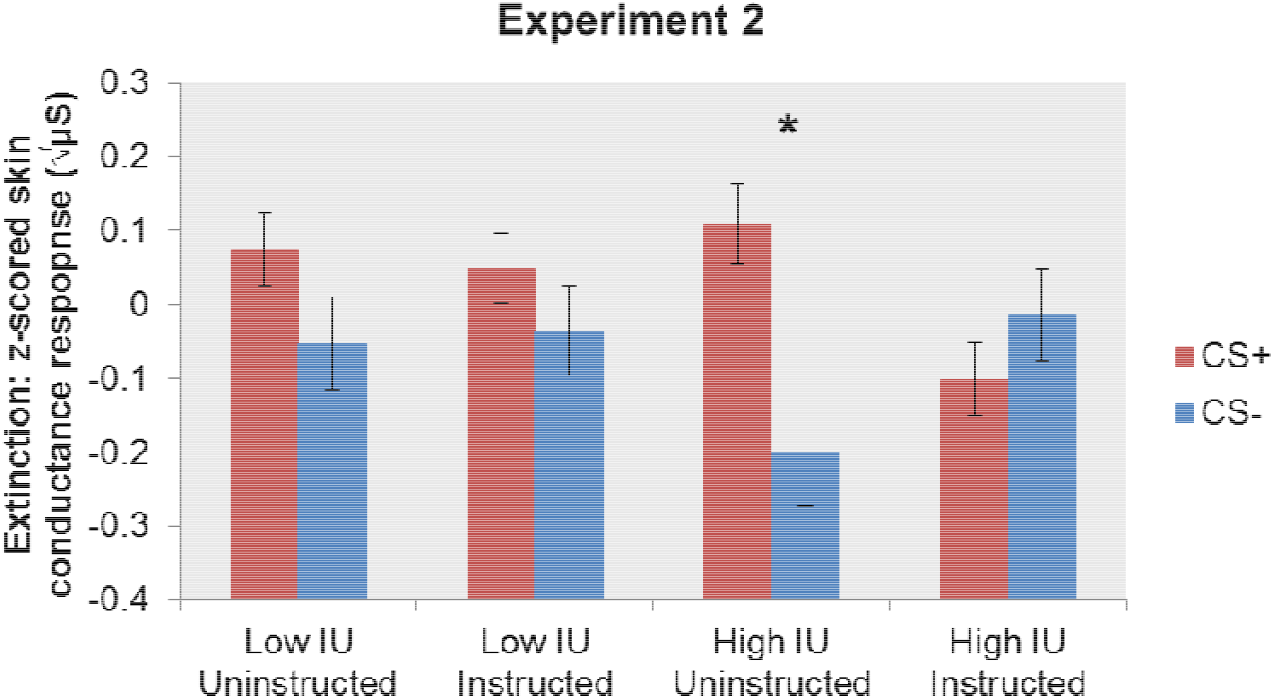
Experiment 2 SCR magnitude results for IU group (controlling for STAI) during threat extinction. Replicating results from experiment 1, only the high IU uninstructed group were found to show differential skin conductance responding to the CS+ versus CS− cue during threat extinction. Bars represent standard error. Square root transformed and z-scored SCR magnitude (μS), skin conductance magnitude measured in microSiemens. Note that the z-scoring was performed within-subjects, across both phases, thus explaining the negative values for a number of conditions.

## Experiment 2: Conclusion

The majority of the results from experiment 2 were similar to experiment 1. As in experiment 1, we observed a similar pattern of acquisition on the SCR magnitudes and expectancy ratings. However, the SCR magnitude difference for the CS+ vs. CS− during acquisition was not as strong. For extinction the SCR magnitudes and expectancy ratings were larger for the CS+ vs. CS. Again, during extinction, only the uninstructed high IU group displayed larger SCR magnitudes to the CS+ vs. CS. The other three groups showed no differential SCR magnitudes between the CS+ vs. CS, indicative of extinction. These effects were found irrespective of time (early vs late). We observed no IU differences on the ratings.

## General Discussion

In two experiments, we show that reducing uncertain threat via contingency information promotes threat extinction in high IU individuals, indexed by lessened differential SCR magnitude responding to learned threat vs. safety cues. These results provide further evidence that uncertainty plays a critical role in threat extinction, which may have important implications for current and future anxiety disorder diagnosis and treatment targets.

For both experiments we observed typical patterns of acquisition, where larger SCR magnitudes and expectancy ratings were found for the learned threat vs. safety cues. In both experiments individual differences in IU predicted the extent of extinction. As expected, the uninstructed high IU group’s displayed reduced threat extinction, as shown by larger differential SCR magnitude responding to learned threat vs. safety cues. This result sits alongside previous work, where high IU has been found to be associated with poorer extinction outcomes (Dunsmoor, Campese, Ceceli, LeDoux, & Phelps, 2015; Lucas, Luck, & Lipp, 2018; Morriss, Christakou, & van Reekum, 2015, 2016; Morriss, Macdonald, & van Reekum, 2016). Importantly, the high IU instructed displayed threat extinction, as shown by lessened differential SCR magnitude responding to learned threat vs. safety cues, similar to the low IU groups. The observed IU-related effects on SCR magnitude during extinction for both experiments were specific to IU, over STAI.

The results above suggest that it is the uncertainty during threat extinction that maintains the conditioned response in high IU individuals. This understanding is in line with the modern definition of IU, i.e. ‘IU is an individual’s dispositional incapacity to endure the aversive response triggered by the perceived absence of salient, key, or sufficient information, and sustained by the associated perception of uncertainty’ (Carleton, 2016b, p. 31). Notably, in the current experiment, we provided participants with information for both the learned threat and safety cue. Therefore, we cannot deduce whether it is the uncertainty of the learned threat cue (US omission) or the uncertainty of both the learned threat and safety cue. To tease this apart further, the next step would be to include instructed groups with partial information about the learned threat cue and safety cue separately. We would predict that high IU individuals would show the poorest extinction outcomes for more uncertain versus certain contexts (e.g.: no information, partial information for the learned safety cue, partial information for the learned threat cue and full information would be associated with better extinction for high IU respectively).

Threat extinction learning principles underlie current exposure-based therapies. We can speculate from the current findings that IU may be one of the reasons why some individuals may take longer to benefit from exposure therapies or may be unresponsive to exposure therapies altogether. The results from the current study are promising, as it suggests that high IU individuals are able to use contingency information to alter their behaviour during extinction. Notably, always using or seeking such information to reduce uncertainty is not necessarily a helpful strategy. Indeed, relying on information to reduce uncertainty may be a safety behaviour. However, there may be other types of information high IU individuals can use to help them tolerate uncertainty (e.g. putting more weight on information that leads to positive outcomes) . It will be important to conduct future research with a focus on developing experimental and clinical interventions that use other types of information to speed up or prolong extinction in high IU individuals across disorders with an anxiety component (Craske, Treanor, Conway, Zbozinek, & Vervliet, 2014; Knowles & Olatunji, 2018).

In the current experiments we did not observe time-based effects of IU and threat extinction as we did in our original experiments (Morriss, Christakou, & van Reekum, 2015, 2016). The difference between these experimental findings may be due to the reinforcement rate and timing of the CS. In this study we used a 50% reinforcement rate during the acquisition phase, whilst in our original experiments the rate was 100%. We used a 50% reinforcement rate in part to assess the conditioned response without the potential confound of the sound and to maintain the effect of conditioning during extinction (Leonard, 1975; Livneh & Paz, 2012). In addition, the experiments reported here used a CS of 4 seconds, whilst in our original experiments the CS was 1.5 seconds. From a methodological standpoint, it is advantageous to use a CS with a longer duration as it allows for more SCRs to be captured across all trials. Despite these design differences, IU-related effects were still observed in extinction.

For both experiments the IU-related results in extinction were consistent for SCR magnitude. The majority of research examining the effects of IU on threat conditioning have found significant relationships between IU and psychophysiological measures such as startle and skin conductance (Chin, Nelson, Jackson, & Hajcak, 2016; Morriss, Christakou, & Van Reekum, 2015, 2016; Morriss, Macdonald, & van Reekum, 2016; Morriss, McSorley, & van Reekum, 2017; Sjouwerman, Scharfenort, & Lonsdorf, 2017). For the ratings we observed results with STAI over IU in experiment one for the extinction phase. In experiment two, neither IU nor STAI significantly predicted the expectancy ratings during extinction. To our knowledge only a few studies have observed IU effects on ratings (Morriss, Macdonald, & van Reekum, 2016; Sjouwerman, Scharfenort, & Lonsdorf, 2017). We therefore think that IU may be a more suitable predictor of bodily responses during threat extinction. The lack of consistent patterns between psychophysiological and rating measures for IU may also be due to the time between phasic cue events and rating periods in the experiments, where ratings are provided retrospectively.

To improve the generalisability of results future studies should aim to replicate IU and extinction effects in more diverse samples (see Supplementary Material on undergraduate psychology sample). It may be of interest to examine whether the current results are similar to clinical samples with high IU. The mean IU score in the current sample was: (1) approx. 10 points higher than those reported in student samples from North America (Carleton, Norton, & Asmundson, 2007), and (2) approx. 7 points above the clinical cut-off used for patients with GAD (Dugas & Ladouceur, 2000). Hence, findings obtained from the samples in this study likely have relevance for clinical research.

In conclusion, these initial results provide insight into how uncertainty during threat extinction may maintain the conditioned response in high IU individuals, which will be relevant for understanding uncertainty-induced anxiety diagnostics and treatment targets (Carleton, 2016a, 2016b; Grupe & Nitschke, 2013). Further research is needed to explore how individual differences in IU modulate learned associations during extinction with and without instruction, and across longer time frames in the laboratory and clinic.

## Acknowledgements

This research was supported by the Centre for Integrative Neuroscience and Neurodynamics (CINN) at the University of Reading. The authors wish to thank Megan James, Francesco Saldarini, Maria Petridou, Tianyao Tong, Kate Cullen, Sophie Szymkowiak, and Maria Dimciu for their help in data collection. The authors thank the participants who took part in this study. The authors received no funding from an external source. To access the data, please contact Dr. Jayne Morriss.

## Footnotes

1 To assess whether the results during threat extinction were due to IU and not STAI, we conducted the same analysis with groups split by instruction and STAI. The instructed and uninstructed groups were matched on STAI: low STAI uninstructed (*n* = 18, *M* = 34.5; *SD* = 5.09); low STAI instructed (*n* = 14, *M* = 32.00; *SD* = 5.09); high STAI uninstructed (*n* = 10, *M* = 50.60; *SD* = 5.13); high STAI instructed (*n* = 15, *M* = 52.93; *SD* = 4.61). No significant interactions with STAI group were found, max *F* = 1.114.

2 To check that the results during threat extinction were due to IU and not STAI, we conducted the same analysis with groups split by instruction and STAI. The instructed and uninstructed groups were matched on STAI: low STAI uninstructed (*n* = 22, *M* = 36.09; *SD* = 4.72); low STAI instructed (*n* = 20, *M* = 36.75; *SD* = 3.87); high STAI uninstructed (*n* = 18, *M* = 53.00; *SD* = 6.35); high STAI instructed (*n* = 19, *M* = 50.53; *SD* = 5.35). No significant interactions with STAI group were found, max *F* = 1.509.

